# Neurons alter endoplasmic reticulum exit sites to accommodate dendritic arbor size

**DOI:** 10.1101/2022.11.03.515099

**Authors:** Ruben Land, Richard Fetter, Xing Liang, Christopher P. Tzeng, Caitlin Taylor, Kang Shen

## Abstract

Nervous systems exhibit dramatic diversity in cell morphology and size. How neurons regulate their biosynthetic and secretory machinery to support such diversity is not well understood. Endoplasmic reticulum exit sites (ERESs) are essential for maintaining secretory flux, and are required for normal dendrite development, but how neurons of different size regulate secretory capacity remains unknown. In *C. elegans, we* find that the ERES number is strongly correlated with the size of a neuron’s dendritic arbor. The elaborately branched sensory neuron, PVD, has especially high ERES numbers. Asymmetric cell division provides PVD with a large initial cell size critical for rapid establishment of PVD’s high ERES number before neurite outgrowth, and these ERESs are maintained throughout development. Maintenance of ERES number requires the cell fate transcription factor MEC-3, *C. elegans* TOR (*ceTOR/let-363*), and nutrient availability, with *mec-3* and *ceTOR/let-363* mutant PVDs both displaying reductions in ERES number, soma size, and dendrite size. Notably, *mec-3* mutant animals exhibit reduced expression of a *ceTOR/let-363* reporter in PVD, and starvation reduces ERES number and somato-dendritic size in a manner genetically redundant with *ceTOR/let-363* perturbation. Our data suggest that both asymmetric cell division and nutrient sensing pathways regulate secretory capacities to support elaborate dendritic arbors.

## Introduction

Neurons exhibit a large diversity in morphology and size. A single nervous system contains small neurons with simple morphology and very large neurons with extremely complex neurites. How neurons adjust their biosynthetic and secretory activity to match their morphology and size is not well understood. It is well known that cell types with secretory functions such as pancreatic acinar cells, have expanded rough ER and Golgi apparatus (Gorelick and Jamieson, 2012). Work in secretory cells and mammalian cell lines demonstrates that early secretory organelle size and number is associated with changes in secretory cargo load (Clermont et al., 1993; Farhan et al., 2008a; Griffiths et al., 1985). However, it is not currently understood how post-mitotic neurons of different types regulate secretory capacity to accommodate divergent cell sizes *in vivo*.

Examination of endoplasmic reticulum exit sites (ERESs) may offer a valuable proxy for gauging secretory capacity in neurons *in vivo*. ERESs export correctly folded proteins from the Endoplasmic Reticulum (ER), via active cargo selection and bulk flow through a system of vesicles or tubules, to subsequent secretory compartments such as the Golgi (Barlowe and Helenius, 2016; Peotter et al., 2019; Weigel et al., 2021). As the first step in the secretory pathway, ERESs are intimately linked to a cell’s overall secretory flux (Barlowe and Helenius, 2016). Indeed, ERES size and number have been shown to increase with elevated secretory load (Farhan et al., 2008a). Specialized cell types with high biosynthetic and secretory capacities can display dozens to hundreds of ERESs, while simpler cells can display as few as a single ERES (Saegusa et al., 2022; Stephens, 2003; Warren, 2013; Yelinek et al., 2009). In *Drosophila* neurons, genes involved in ERES function have been shown to play an important role in dendritic arbor development (Ye et al., 2007).

The worm, *Caenorhabditis elegans*, offers an attractive system for studying how neurons regulate secretory capacity to support diverse cell size. *C. elegans* hermaphrodites have 302 neurons of widely varying size and complexity, many of which can be genetically manipulated on the individual cell level. In addition, the transparent *C. elegans* allows for visualization of secretory organelle formation and dynamics within a single neuron throughout its development. The vast majority of *C. elegans* neurons a have simple cell morphology with 1 or 2 unbranched neurites (White et al., 1986). However, the multimodal sensory neuron PVD develops an expansive dendritic arbor with over 150 terminal branches that span nearly the entire length of the worm (Oren-Suissa et al., 2010; Smith et al., 2010). How PVD adapts the function or structure of its secretory pathway to accommodate the secretory flux required to establish such a large dendrite is not known.

In this study, we determined ERES number in a subset of *C. elegans* neurons of different sizes and morphological complexities. We report that neurons with larger dendrites had significantly higher ERES numbers in their somas, with PVD displaying the highest number of ERESs. Our data indicate that asymmetric cell division, resulting in a large initial soma size, establishes PVD’s high ERES number before neurite outgrowth. Subsequent maintenance of ERES number during PVD development is coupled with expansive somato-dentritic growth. Starvation, as well as disruption of the cell fate transcription factor MEC-3 and the master metabolic regulator *C. elegans* TOR (*ceTOR/let-363*), all resulted in a coordinated reduction of ERES number, dendritic and soma size. Moreover, *mec-3* perturbation reduced expression of a *ceTOR/let-363* transcriptional reporter in PVD, and the impact of starvation on ERES number is redundant with that of the *ceTOR/let-363*-deletion in PVD. These data suggest that cell fate transcription factors in PVD upregulate both initial cell size and nutrient sensing pathways, which together establish the high biosynthetic and secretory capacity required to generate and maintain a large dendritic arbor.

## Results

### Larger neurons have more ER-exit sites

To understand the molecular mechanisms that generate the morphological diversity of neurons, we postulated that biosynthetic and membrane trafficking pathways are tuned to neuron size and complexity. To test this hypothesis we visualized ERESs in neuronal types with different morphological complexity, using an endogenous, tissue-specific tag of ERES protein SEC-16 (GFP::FLPon::SEC-16). We observed that neurons with larger, more complex morphologies have significantly higher numbers of ERESs in their cell bodies or somas (Fig. 1). Touch receptor neurons (TRNs) AVM and PVM, each with fewer than 5 terminal neurite branch points, typically have three to four distinct somatic ERESs. In contrast, significantly higher numbers of ERESs are found in the somas of multimodal sensory neurons FLP and PVD, each of which have highly branched dendrites with dozens of terminal neurite branch points (Fig. 1a, b, c). PDE, a smaller sensory neuron within the same lineage as PVD, has fewer ERESs than PVD. Comparing individual FLP and PVD neurons, we also found a trend that cells with higher branch numbers have more ERESs (Fig. 1c). To validate the discrete punctate nature of the fluorescent ERES marker, we reconstructed PVD and PDE cell somas by serial section electron microscopy (EM). The EM images indicated six discrete Golgi stacks in PVD soma, each with one or two associated ER exit sites, corresponding well with results from fluorescence microscopy (Fig. 1d-g). We chose PVD as a system in which to further examine the relationship between ERES function, ERES number and dendrite size, due to its large size, high ERES number, elaborately branched and quantifiable dendrites, and its well-characterized morphogenesis (Zou et al., 2016).

**Fig.1:**
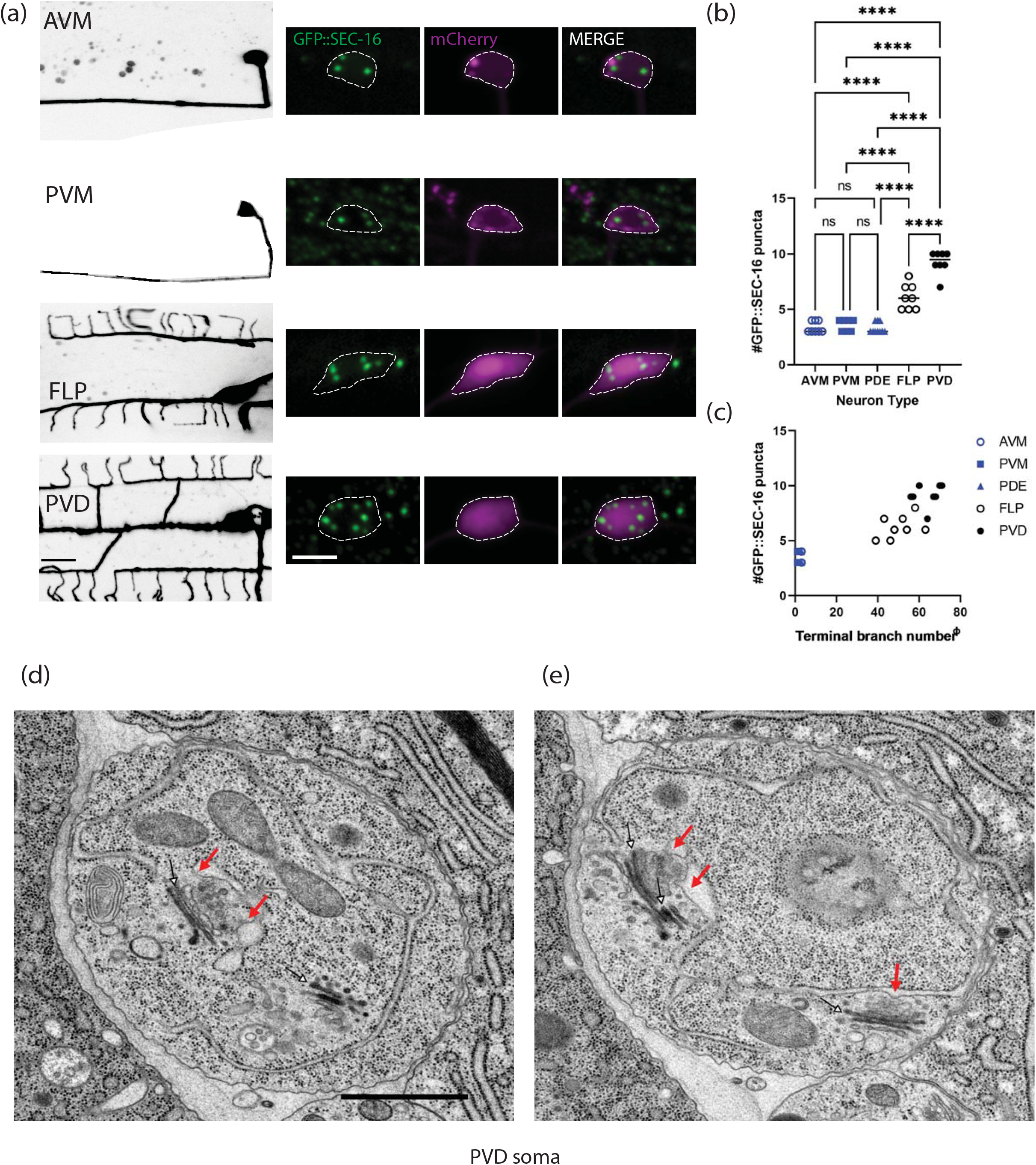
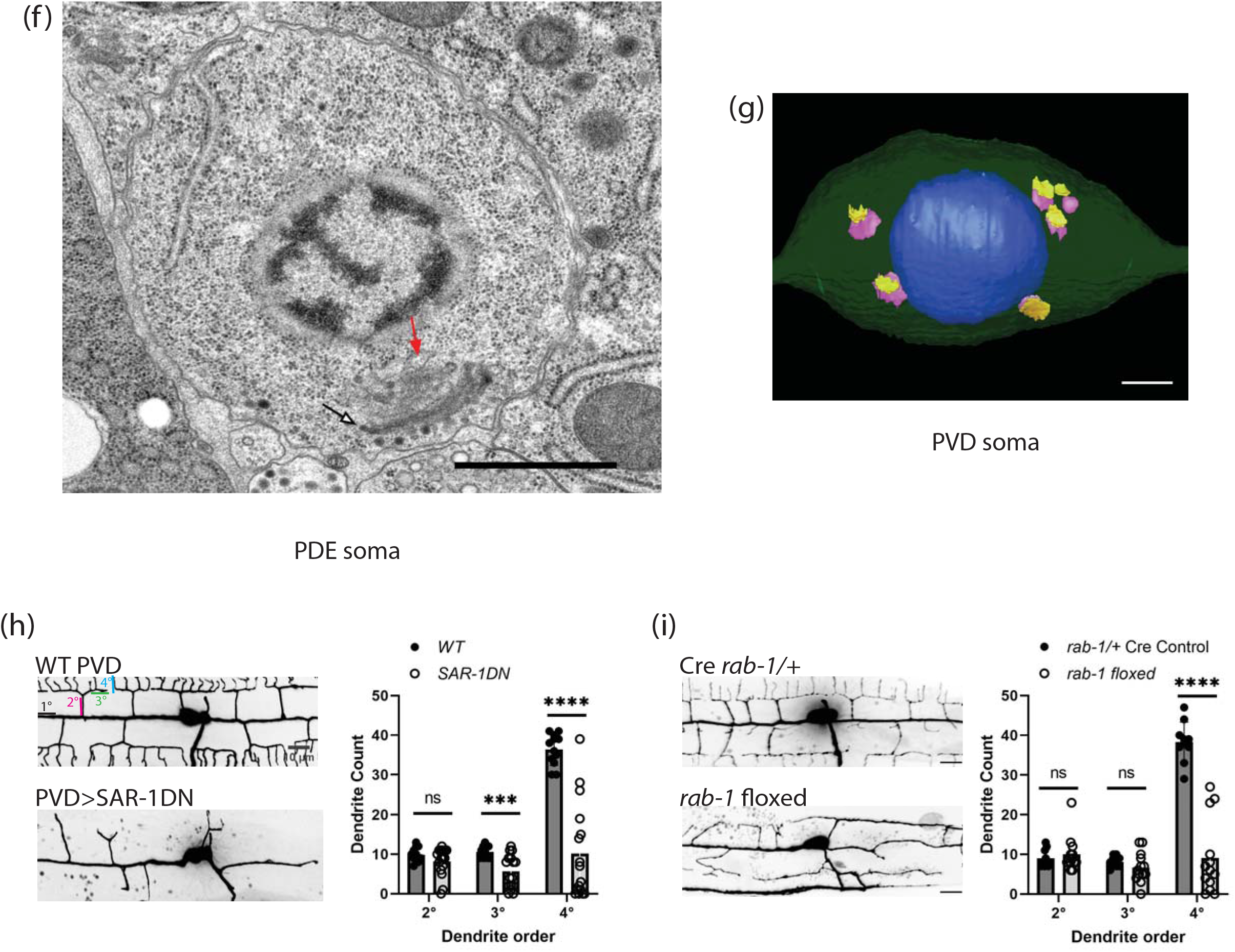
Larger neurons have more ER-exit sites (ERESs) in their somas. (a) Confocal fluorescence microscopic images of neuron morphology on the left (scale bar = 10μm), with closeups of soma and an endogenous ERES marker, GFP::SEC-16, on the right (scale bar = 5μm). Background of PVM image was removed to more clearly show PVM morphology. (b) Quantification of number of GFP::SEC-16 puncta per soma in different cell types. Significance determined by one-way ANOVA, ****p<.0001, n≥7 animals. (c) Quantification of number of GFP::SEC-16 puncta per soma in different cell types vs. number of terminal dendrites (Φ see methods). Pearson correlation between ERES number and terminal dendrite number: r=0.9282, R^2^=0.8615, p<.0001, n=41 neurons with at least 7 of each type. Electron microscopic (EM) images of PVD (d, e) and PDE (f) soma. Red arrows indicate ER-exit sites (ERES). Black & white arrows indicate Golgi stacks. Scale bars =1μm. (g) 3-D model of PVD soma constructed from EM images showing nucleus in blue, with cis (purple) and trans (yellow) cisternae of Golgi. Scale bar =1μm. (h) Confocal fluorescence microscopic images of WT PVD and PVD with exogenously expressed SAR-1 dominant negative (SAR-1DN), with quantifications of secondary (2°), tertiary (3°) and quaternary (4°) dendrites in WT (n=12 animals) and SAR-1-DN (n=16 animals). Significance determined by multiple t tests, ***p<.001, ****p<.0001. (i) Confocal fluorescence microscopic images of Cre *rab-1*/+ heterozygous control PVD (n=10) and *rab-1* floxed PVD (n=13), with corresponding dendrite quantifications as in (h). Results were additionally repeated in at least two independent experiments.

### ERES function is required for PVD dendrite size

To test the relationship between ERES function and dendrite size directly, we disrupted ERES function in PVD, whose elaborately branched dendrite allows for quantitative assessment of dendritic growth defects. PVD dendrite was dramatically reduced as a result of disrupting ERES function via two independent approaches (Fig. 1). First, we expressed a dominant negative version of the ERES protein SAR-1 (SAR-1DN) specifically in PVD. SAR-1 is a small GTPase required for protein trafficking from ER to Golgi(Long et al., 2010; Nakano and Muramatsu, 1989). Overexpression of SAR-1DN severely disrupts dendrite growth in PVD (Fig. 1h). Second, we made a tissue-specific PVD knockout of *rab-1*, which is required for ER to Golgi trafficking (Balklava et al., 2007; Sannerud et al., 2006; Sato et al., 2006). Similar to SAR-1-DN over-expression PVDs, *rab-1*-floxed PVDs exhibited severe branching defects (Fig. 1i), suggesting that ERES function is critical for dendrite growth and branching.

### Dendrite growth is not a prerequisite for increased secretory organelle abundance in PVD

Having established a strong association between ERES function, ERES number and dendrite size, we hypothesized that the number of early secretory organelles might increase in response to rising secretory demand in expanding neurites. We therefore asked whether normal dendrite outgrowth is required for high ERES number in PVD. To test this hypothesis, we disrupted several factors that are critical for PVD dendrite guidance (Dong et al., 2016; Liu and Shen, 2011; Zou et al., 2016). Perturbation of these guidance receptors and regulators including *dma-1, hpo-30* and *kpc-1*, severely disrupts PVD dendrite morphology but does not alter the number of early secretory organelles (Fig. S1). Thus, feedback from dendrite outgrowth and precise arbor shape are not required for high secretory organelle abundance in PVD.

### ERES number is coupled with dendrite size in PVD cell fate mutants

Given that feedback from an expanding dendritic arbor is not required for establishing secretory organelle number, we wondered whether PVD’s high ERES number is instead regulated by cell fate transcription factors that drive PVD dendrite arborization. MEC-3 and UNC-86 are transcription factors that affect many aspects of PVD cell fate including dendrite growth. In *mec-3(e1338)* or *unc-86 (e1416)* mutants, PVD loses nearly all of its dendrites (Fig. 2a, Smith et al., 2013, 2010; Tsalik and Hobert, 2003), and we find a corresponding reduction of ERES number in mutant PVD somas. We also tested whether cell fate perturbations that increase dendritic size and complexity would result in elevated ERES numbers. Wild-type (WT) AVM has a simple neurite morphology. In transcription factor *ahr-1(ju145)* mutants, AVM becomes a cell with a large PVD-like dendritic arbor, cAVM (Fig. 2c, Smith et al., 2013). Notably, cAVMs also have much higher ERES numbers than AVM (Fig. 2d). These results indicate that transcription factors controlling dendrite size also specify ERES numbers, perhaps as part of cell fate programming.

**Fig.2:**
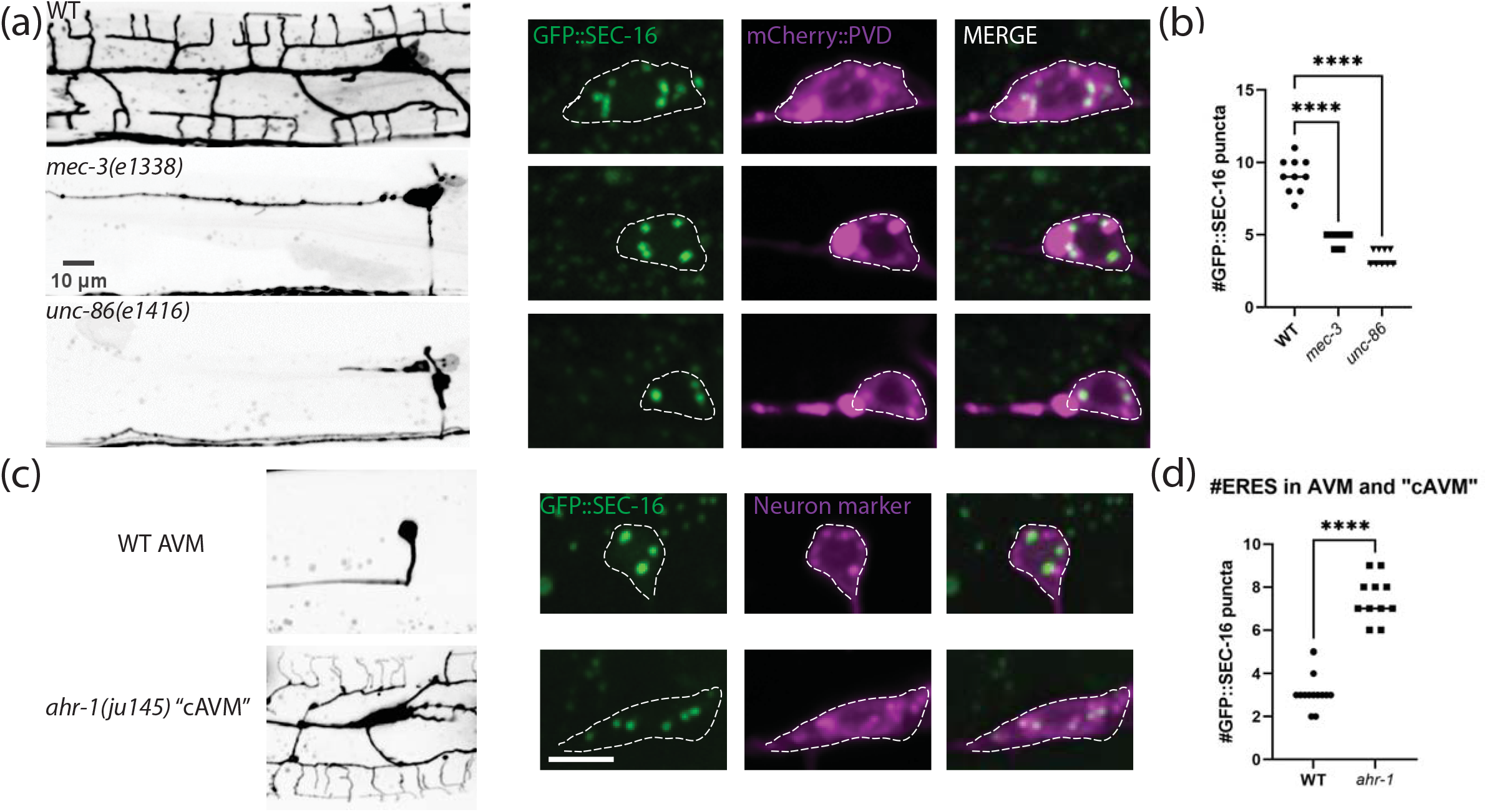
Cell fate mutants that alter neuron size perturb ERES abundance. (a & c) confocal fluorescence microscopy of neuron morphology of wild type (WT) and mutant worms on the left (scale bars = 10μm, with closeups of soma and an endogenous ERES marker, GFP::SEC-16, on the right (scale bar = 5μm). (b & d) Quantification of number of GFP::SEC-16 puncta per soma. Significance determined by one-way ANOVA (n≥9 animals) in (b), and Student’s t-test two-tailed in (d), ****p<.0001, n≥11. Results were additionally repeated in at least two independent experiments.

### ERES number is established early after PVD birth and maintained throughout PVD development

Given that secretory organelle number in PVD is dependent on cell fate transcription factors but independent of dendrite growth, we predicted that ERES number is established early in PVD development. To test this prediction, we determined when ERES number is established, and how ERES number changes during neuron growth by conducting time-lapse recordings and time course experiments throughout PVD birth and development (Fig. 3). These dual-color timelapse recordings of live animals allowed us to observe endogenous SEC-16 dynamics during PVD birth, and track ERES formation and numbers simultaneously with PVD morphology changes. We observed that high ERES number is established within the first 30 minutes of PVD birth, even before neurite outgrowth, and is subsequently maintained throughout PVD development (Fig. 3a, b, c). This rapid establishment of high ERES numbers is consistent with the idea that ERES number is determined via cell fate programming rather than feedback from expanding neurites.

**Fig. 3:**
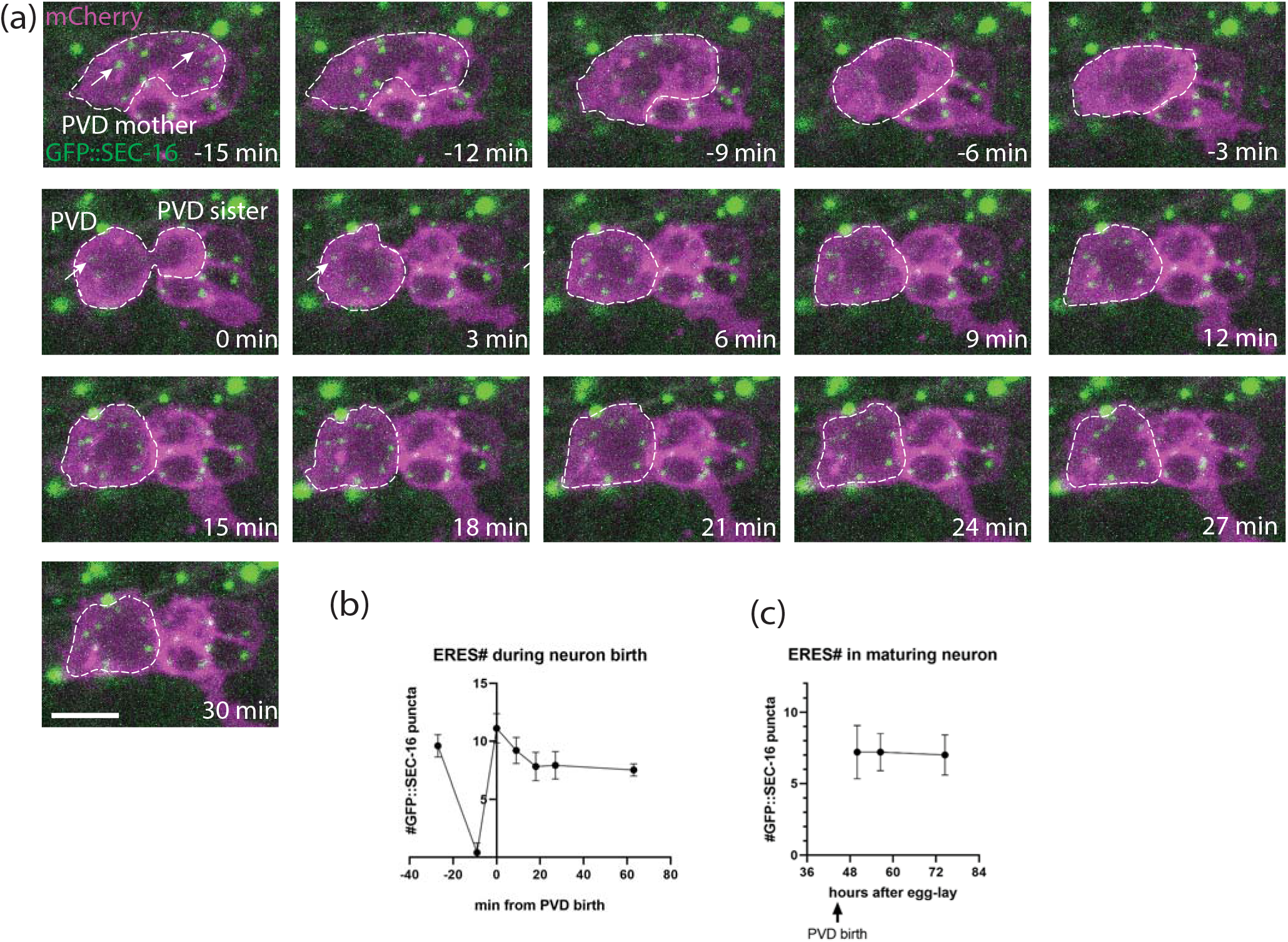
ERES number is established rapidly (Phase 1) after neuron birth and persists over time. (a) Time-lapse confocal fluorescence microscopic images of PVD cell birth in wild-type L2 worms with Plin-32::mCherry::PH labeling neuron membranes and an endogenous fusion protein GFP::SEC-16 labeling ERESs (white arrows). Scale bar = 5μm. (b) Quantification of ERES number from long-term time lapse imaging during PVD birth, n≥9 for each time point. (c) Quantification of ERES number after PVD birth, via confocal microscopy, as worm matures, n=10 for each time point. Results were additionally repeated in at least two independent experiments.

In addition to pinpointing ERES emergence to a narrow time window shortly after PVD birth, our timelapse analyses identified two distinct phases of ERES dynamics after PVD birth that might contribute to final ERES number in mature PVD: ERES establishment and maintenance. The establishment phase begins in the final stages of mitosis, when GFP::SEC-16 starts to coalesce rapidly into puncta that are clearly visible within three minutes of cytokinesis (Fig. 3a). These puncta undergo a brief period of consolidation within the first several minutes of birth, establishing a high number of ERESs before neurite outgrowth (Fig. 3a, b). Subsequently, during the maintenance phase, ERES number remains at consistently high levels throughout PVD development (Fig. 3c). Two central questions emerging from these observations: how high ERES numbers in PVD are established, and how they are they maintained throughout PVD development.

### Cell size at birth impacts ERES number in PVD

Because of the asymmetric cell division of PVDmother observed in Fig. 3a, we hypothesized that large cell size at birth drives establishment of high ERES number. Consistent with previous findings, our timelapse imaging showed that PVD is born significantly larger than its sister cell, PVDsis (Fig. 3a, Fig. 4a, b upper left), which is eventually eliminated via apoptosis (Teuliere et al., 2018) (data not shown). We hypothesized that this asymmetric cell division is required for unequal distribution of cytoplasm, membranes, and ERES components, and may be responsible for seeding the high ERES number in PVD soon after birth. Consistent with this idea, the initial ERES number in PVD after consolidation is consistently higher than the ERES number in the smaller PVDsis (Fig. 4a). To further test the impact of initial PVD size on ERES number, we examined initial ERES number under conditions in which asymmetric cell division was disrupted. In *unc-86(e1416)* mutants, PVD and PVDsis have similar cell size. We found that *unc-86(e1416)* mutant PVD and PVDsis showed similar ERES numbers, which fall in between the values of WT PVD and WT PVDsis (Fig. 3b and top-right, 3c). These results are consistent with the notion that cells that are born larger, form more ERESs. However, *unc-86(e1416)* mutants exhibit abnormalities in asymmetric cell division and PVD cell fate pathways, both of which likely impact ERES number. It thus remained unclear whether the ERES phenotype in the *unc-86(e1416)* mutants is due to defects in asymmetric cell division or unrelated cell fate pathways.

**Fig. 4:**
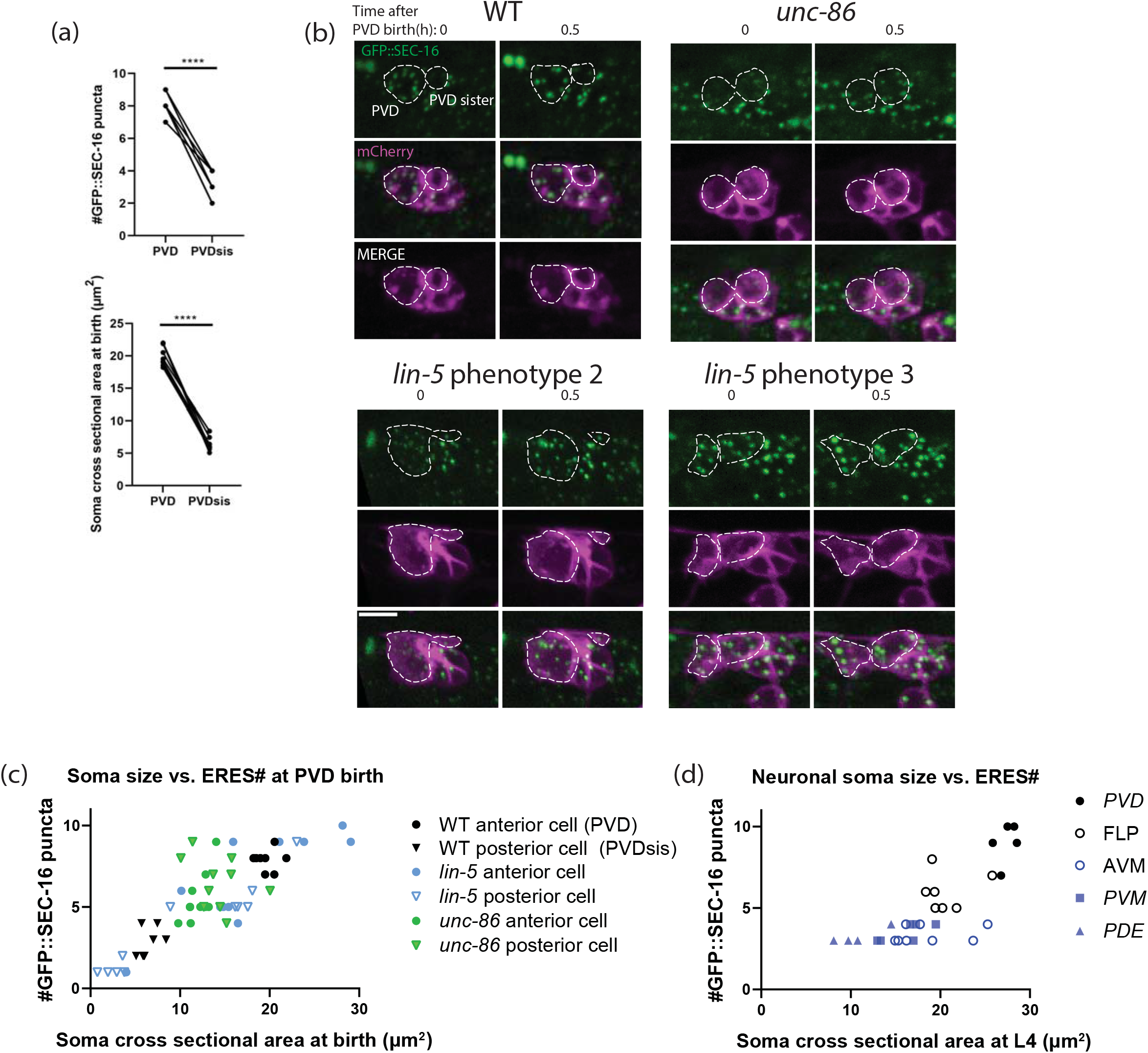
Asymmetric cell division is important for establishment phase of ERES number. (a) Quantification of ERES number shortly after birth (top), and soma cross-sectional area at birth (bottom) in PVD and PVDsister. Significance determined by Student’s t-test two-tailed, ****p<.0001, n=8 for each cell type. (b) Representative images of wild type (WT) and mutant worms at PVD birth and 0.5 hour after birth. *lin-5* mutants sometimes increase asymmetry of division (bottom left) and sometimes reduce asymmetry of division (bottom right). Scale bar = 5μm. (c) Quantification of cross-sectional soma area at birth vs. ERES number 30 minutes after birth of PVD and PVDsister in WT and mutants. Pearson correlation between soma size and ERES number: r=0.8404, R^2^=0.7062, p<.0001, n=58 cells from 29 animals (WT: n=8, *lin-5*: n=11, *unc-86*: n=10). (d) Quantification of cross-sectional soma area vs. ERES number across mature neuron types at larval stage L4. Pearson correlation between soma size and ERES number: r=0.8395, R^2^=0.7048, p<.0001, n=34 cells with at least 6 of each type. Results were additionally repeated in at least two independent experiments.

The impact of cell size on establishment of ERES number was directly tested using a *lin-5* mutation. LIN-5/NUMA directly regulates asymmetric cell division and cell size by specifying spindle position during mitosis without directly involving transcription factors (Colombo et al., 2003; Gotta et al., 2003; Srinivasan et al., 2003). It has been shown that *lin-5* mutations alter relative daughter cell size without changing anterior-posterior polarity (Jankele et al., 2021). To bypass the lethality of *lin-5* null mutants, we generated a conditional *lin-5* strain by tagging the endogenous *lin-5* locus with a zf ubiquitination sequence (Armenti et al., 2014). We used the *unc-86* promoter to express the ubiquitin ligase *zif-1* to degrade zf-tagged LIN-5 in PVDmother shortly before PVD birth (Finney and Ruvkun, 1990). This manipulation displaces the spindle during PVDmother division, resulting in several categories of cell division phenotypes that differ from animal to animal: 1) Failed cell division; 2) exaggerated asymmetry of daughter cells where the size difference between the daughter cells is larger than normal (Fig. 4b lower left); 3) reduced asymmetry of daughter cells, where the daughter cells display a less pronounced size difference (Fig. 4b lower right), and 4) WT-like asymmetry of daughter cells. We took advantage of these different phenotypes to compare initial ERES numbers in daughter cells of differing soma size at birth. We found a strong correlation between initial soma size and initial ERES number (Fig. 4c), suggesting that asymmetric cell division and large cell size at birth drive the establishment phase of high ERES numbers in PVD. Notably, we also observed a strong correlation between soma size and ERES number in mature neurons of different types (Fig. 4d). Together with Figure 1c, this indicates that neurons of increasing dendrite size have larger somas and higher ERES numbers.

### PVD cell fate transcription factor MEC-3 is required for ERES number maintenance but not establishment

Having elucidated a mechanism for the establishment phase of high ERES number, we investigated mechanisms underlying the maintenance phase of ERES number during PVD development. Because MEC-3 acts downstream of UNC-86 (Chalfie and Au, 1989; Xue et al., 1993, 1992), and the mature PVD neuron in the *mec-3(e1338)*-mutant has a less severe phenotype than *unc-86(e1416)* (Fig. 2b), we hypothesized that the *mec-3(e1338)* mutation would preserve asymmetric cell division, but disrupt the later maintenance of ERES number. Consistent with this hypothesis, in *mec-3(e1338)* mutants, asymmetric cell division proceeds as normal, and we observe that the initial ERES number is similar to that of WT (Fig. 5a, b left). However, by the L4 stage, *mec-3(e1338)* mutants showed dramatically reduced number of ERESs (Fig. 2a, b, Fig. 5a), indicating that ERES number is actively maintained in a MEC-3-dependent manner as PVD develops. Indeed, time course analysis of *mec-3(e1338)* mutants show that ERES numbers in *mec-3(e1338)* are established at WT levels shortly after PVD birth, but gradually decline as the primary dendrite grows and the worm develops (Fig. 5a, b). Given the connection between soma size and initial ERES number observed in Figure 4, we wanted to test if *mec-3(e1338)* PVD soma size differs from WT. We observed that, like ERES number, initial soma size in WT and *mec-3(e1338)* does not significantly differ (Fig. 5b left). However, by L4, *mec-3(e1338)* PVD has a significantly smaller soma size than WT PVD (fig. 5b right), further supporting an association between ERES number and soma size. While WT soma size increases as PVD develops, *mec-3(e1338)* soma size lacks that increase (Fig. 5b). Notably, in *unc-86(e1416)* animals, where both asymmetric cell division and cell fate pathways are disrupted in PVD, we see a faster, more severe reduction in ERES number over time than in *mec-3(e1338)* mutants (Fig. 5a). A reasonable interpretation of this observation is that both ERES establishment via asymmetric cell division and ERES maintenance via cell fate pathways are required for ERES number in the mature PVD neuron.

**Fig. 5:**
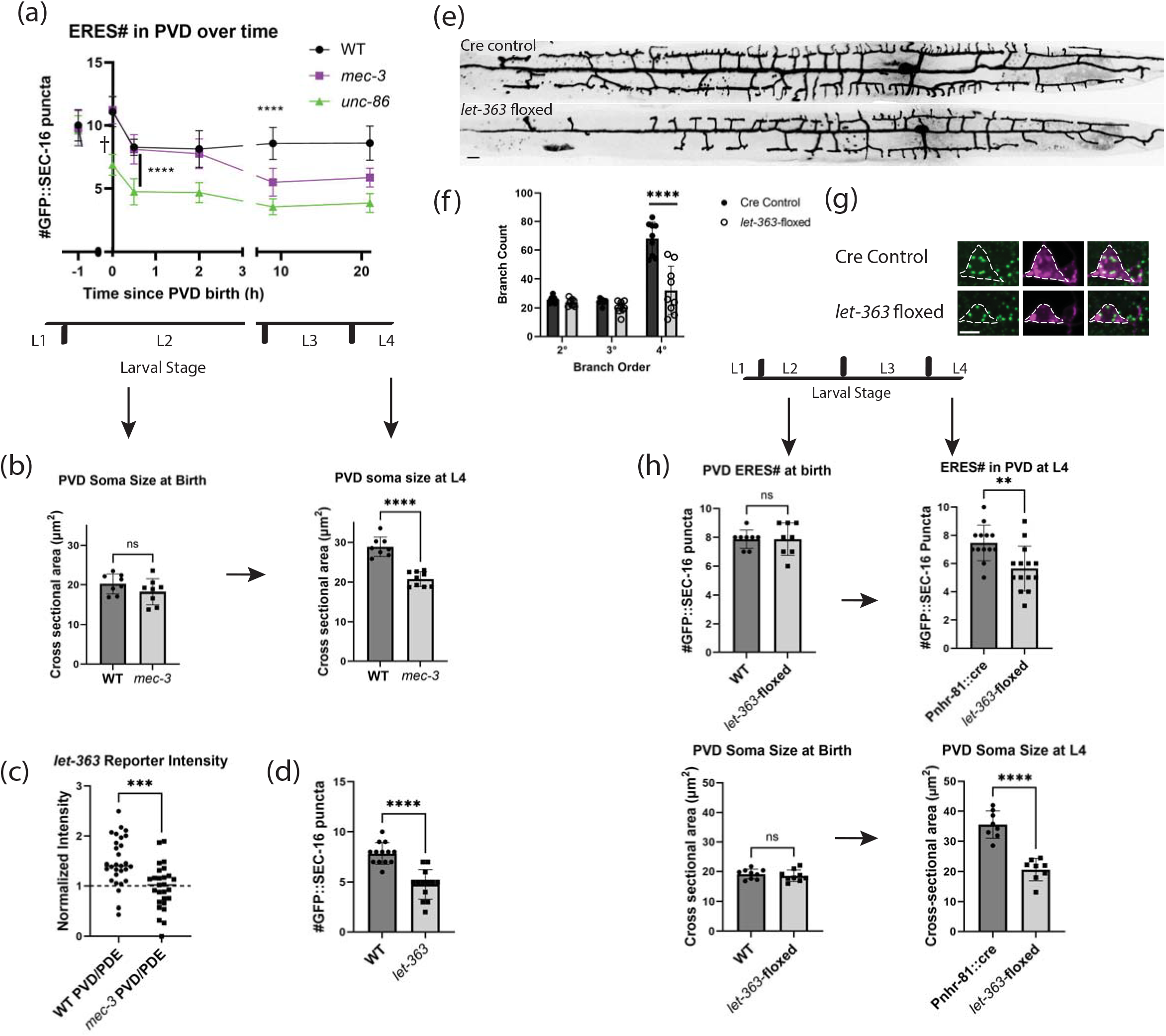
MEC-3 and LET-363 maintain ERESs during neurite outgrowth in PVD. (a) Time course of ERES number in wild type (WT) and cell fate mutants. †Points at t=0 (PVD birth) were counted from time-lapse images to provide accurate timing information. Significance determined by one-way ANOVA, ****p<.0001, n≥8 animals for each genotype at each time point. (b) Soma cross-sectional area of WT and *mec-3* PVD at birth (left) and larval stage 4 (right). Significance determined by Student’s t test two-tailed ****p<.0001, n≥8 animals. (c) P*let-363*::GFP ceTOR transcriptional reporter intensity in PVD normalized to PDE intensity in WT and *mec-3* animals at larval stage 3. Significance determined by unpaired Student’s t test two-tailed ***p<.001, n=29 (WT) & 26 (*mec-3*) animals. Data is from two independent experiments. (d) Quantification of number of GFP::SEC-16 puncta in wild type vs. ce*TOR/let-363(ok3018)* constitutive mutant animal PVD somas. Significance determined by Student’s t test two-tailed ****p<.0001, n=13 for each genotype. (e) Confocal fluorescence microscopic images of PVD morphology in control and mutant worms (scale bar = 10μm), with (f) quantification of branches. Significance determined by multiple t tests ****p<.0001. (g) Confocal fluorescence microscopic images of PVD soma and endogenous ERES marker, GFP::SEC-16 in Pnhr-81::cre control worms (top) and conditionally floxed *let-363* worms (bottom). Scale bar = 5μm. (h) Quantification of number of GFP::SEC-16 puncta (top) and soma cross-sectional area (bottom) per PVD soma in control and mutant L2 (left) and L4 (right) worms. Significance determined by Student’s t test two tailed, **p<.01, ****p<.0001, n≥8 animals. Larval stages from youngest to oldest: L1, L2, L3, L4. YA = Young Adult. Results were additionally repeated in at least one (c & h left) or two independent experiments.

### ceTOR/LET-363 expression is enhanced in PVD by MEC-3

Given the overall lack of somato-dendritic growth in *mec-3(e1338)* mutant PVD, we wondered whether MEC-3 might upregulate expression of master growth regulators. In the vertebrate, the master metabolic regulator mTOR has been associated with overgrowth of neuronal somas in specific contexts (Kwon et al., 2003, 2001). TOR signaling has also been associated with cell size in a variety of cell types (Gonzalez and Rallis, 2017). We hypothesized that MEC-3 promotes expression or activity of *C. elegans* TOR (ceTOR) in PVD to promote increased biosynthetic and secretory capacity, and thus, ERES number in PVD. To test this hypothesis, we first examined a transcriptional reporter of *let-363* developed by Long et al. (Long et al., 2002). In order to control for expression level differences between animals, we normalized *ceTOR/let-363* reporter levels in PVD within each animal to reporter levels in PDE, which does not express MEC-3 (Way and Chalfie, 1989). We found that normalized reporter levels in WT PVD were significantly higher than in *mec-3(e1338)* (Fig. 5c). These results suggest that MEC-3 enhances ceTOR/LET-363 expression in PVD.

### ceTOR/LET-363 is required for maintenance of ERES number and somato-dendritic growth in PVD

Given TOR’s well-known role in promoting biosynthesis (Keith Blackwell et al., 2019; Laplante and Sabatini, 2009; Nandagopal and Roux, 2015), we wondered whether upregulation of ceTOR/LET-363 by MEC-3 in PVD might be responsible for PVD’s high ERES number and/or expansive somato-dendritic growth. First, we asked if *let-363* activity is required for PVD’s high ERES number. In the constitutive *let-363(ok3018)* mutant, which exhibits a general growth arrest of *C. elegans*, we observe a reduction in ERES number in PVD soma (Fig. 5d). To ask if LET-363 is required cell autonomously for ERES number, we built a conditional *let-363* allele using the Cre-lox system to disrupt LET-363 specifically in PVD lineage cells (Fig. 5e, f, g). When *let-363* was deleted, ERES number was normal at PVD birth, but significantly reduced by the time PVD reaches maturity at the L4 larval stage (Fig. 5h). Similarly, the conditional *let-363* mutant PVD also displayed normal soma size at PVD birth but reduced soma size by L4. Additionally, this allele showed a significant reduction in the size of PVD’s dendritic arbor compared to Cre controls (Fig. 5e). The striking similarity between the conditional *ceTOR/let-363* phenotypes and the *mec-3(e1338)* phenotypes argues strongly that ceTOR/LET-363 is part of the MEC-3 dependent mechanisms necessary for PVD maturation.

### Starvation reduces ERES number and growth during critical period of development

Because TOR activity typically requires nutrient signals, and nutrient deprivation has been shown to disrupt ERESs (Zacharogianni et al., 2011), we hypothesized that TOR’s positive impact on ERES number in PVD requires the availability of nutrients (Baugh and Hu, 2020; González and Hall, 2017; Keith Blackwell et al., 2019). To test the effect of nutrient restriction on PVD ERES number, we removed worms from food just after PVD birth. The following day, starved worms displayed significantly fewer PVD ERESs than fed worms (Fig. 6a, b). In addition, PVD soma were significantly smaller in starved vs. fed worms (Fig. 6c). To test whether the TOR-mediated and nutrient-mediated mechanisms on ERES act in the same genetic pathway, we starved WT and *let-363*-floxed worms. We observed that there was no significant additional reduction of PVD ERESs in the *let-363* worms upon starvation (Fig. 6d left). Similarly, there was no significant difference between WT starved worms and *let-363* starved worms (Fig. 6d left). These results are consistent with the notion that the impact of starvation on PVD is mediated at least in part through inactivation of *ceTOR/let-363*. Given our evidence of MEC-3-driven upregulation of LET-363 in PVD, we hypothesized that part of *mec-3(e1338)* mutant’s impact on ERES number in PVD is due to reduced LET-363 activity in these mutants. This hypothesis predicts that inactivation of TOR by starvation of *mec-3(e1338)* mutant animals would only have a small, if any, additional impact on ERES reduction compared to fed *mec-3(e1338)* mutant animals. We tested whether *mec-3(e1338)* and nutrient deprivation have independent effects on ERES number by starving *mec-3(e1338)* and WT worms. As expected, starved *mec-3(e1338)* worms only exhibit a small additional ERES reduction, as compared to WT starved worms and *mec-3(e1338)* fed worms (Fig. 6d right). This is consistent with the idea that ERES number reductions following *mec-3(e1338)* and starvation manipulations are partially but perhaps not fully redundant. Together, these findings suggest that MEC-3 drives a transcriptional program that includes increased *ceTOR/let-363* expression, which helps to maintain PVD’s high ERES numbers in the presence of nutrients. Notably, the reduction of ERES number due to starvation was eliminated in mature starved animals (Fig. 6e), raising the possibility that ERES number stabilizes with age.

**Fig. 6:**
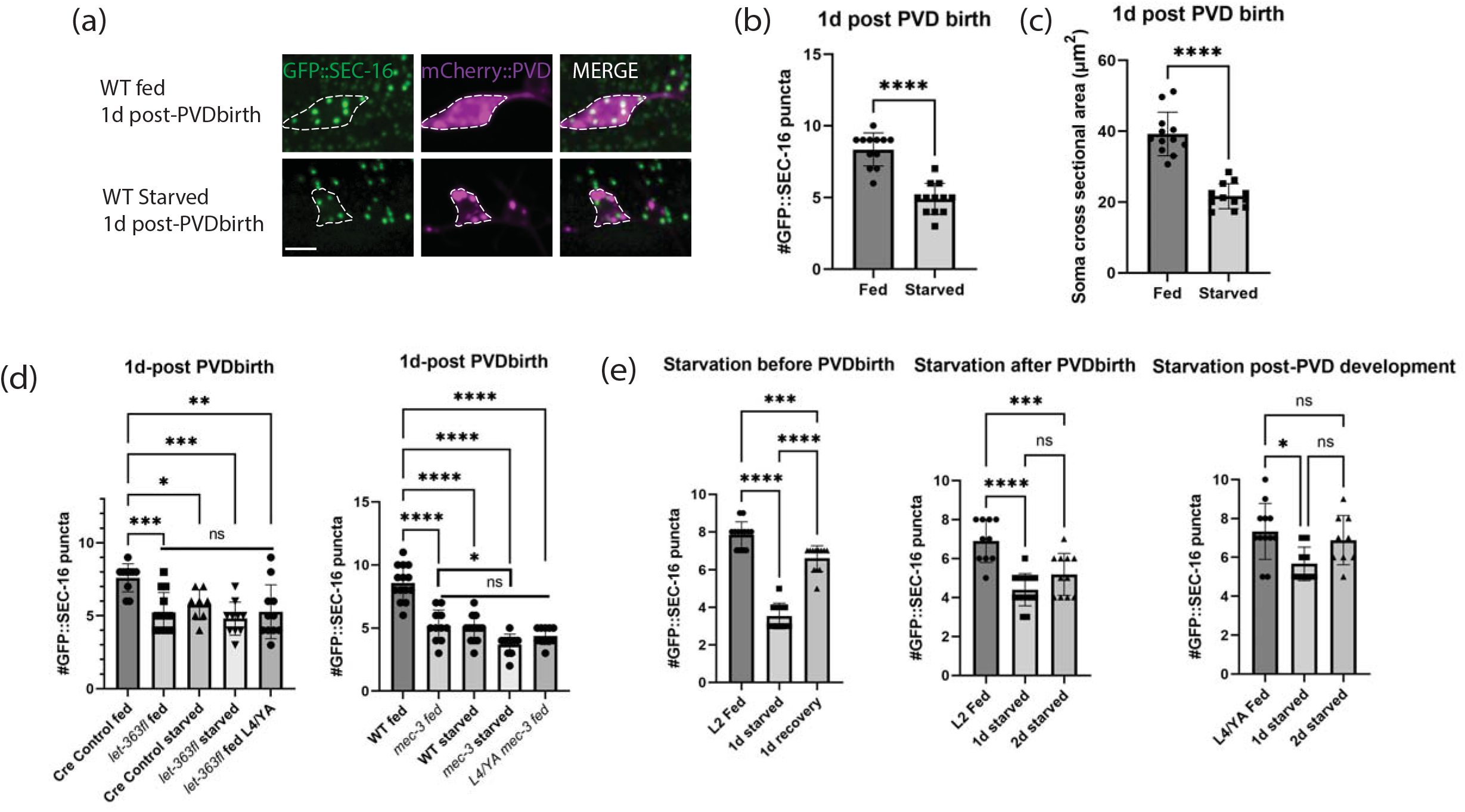
Stage-specific nutrient deprivation reduces PVD ERES number. (a) Confocal fluorescence microscopy close-ups of PVD soma and endogenous ERES marker, GFP::SEC-16, one day after PVD birth in fed or 1-day starved wild type animals (scale bar = 5μm). PVDs in fed and starved animals were quantified for (b) number of PVD GFP::SEC-16 puncta (n=12 per condition), and (c) soma size (n=12 per condition) measured as described in methods. Significance determined by Student’s t-test two-tailed, ****p<.0001. (d) Number of GFP::SEC-16 puncta in PVD soma of fed vs starved animals in Cre-control vs *let-363*-floxed (left, n≥8 animals for each condition) and wild type vs *mec-3* animals (right, n≥10 animals for each condition). Animals were counted one day after PVD birth except where indicated. Fed animals at this time point were in late L3 or early L4 stages. (e) Number of GFP::SEC-16 puncta in wild type animals removed from nutrients at different stages of PVD development, and starved for either one day and recovered (left, n≥11), or two days (middle and right, n≥9). Significance for (d) & (e) determined by one-way ANOVA, *p<.05,**p<.01,***p<.001, ****p<.0001. Results were additionally repeated in at least two independent experiments.

In summary, we show that ERES dynamics are organized into two phases, both of which contribute to high ERES number in PVD. During the establishment phase, UNC-86 and LIN-5 drive asymmetric cell division to generate a large PVD soma at the expense of a small and dying sister cell. Our data suggest that the large soma leads to high ERES number in PVD due to cytoplasmic scaling. During the maintenance phase, as neurites emerge, MEC-3 drives ceTOR/LET-363, which, in the presence of nutrients, maintains high ERES number and promotes growth of the somato-dendritic compartment. Our results suggest a model in which transcription factors, master metabolic regulators and nutrient availability coordinate to specify developmental parameters including soma size, ERES number and dendrite size, which together determine neuronal cell fate.

## Discussion

In this study, we show that ER-exit site (ERES) number is predictive of neuron size and complexity, and identify mechanisms by which large, complex neurons establish and maintain elevated ERES numbers. ERESs export the majority of correctly folded cargo from the ER, and their structure and function is thus critical for maintaining secretory capacity to enable cellular growth and health, particularly in neurons (Barlowe and Helenius, 2016; Peotter et al., 2019; Tang, 2021). We observed a strong correlation between ERES number and dendrite size across cell types and cell fate mutants, with PVD, *C. elegans*’ largest neuron, displaying the highest number of ERESs. ERESs, thus may be regarded as reliable indicators of neuronal biosynthetic and transport activity. We discovered that high ERES number in PVD is established rapidly after neuron birth before neurite outgrowth, and is actively maintained throughout PVD development, using both timelapse imaging and genetic perturbation experiments. Establishment of high ERES number is driven by asymmetric cell division and the resultant large cell size at birth. In turn, maintenance of ERES number is regulated by the transcription factor MEC-3, nutrient availability, and *C. elegans* TOR (*ceTOR/let-363*), a major nutrient sensor and regulator of cell growth. Our data support the notion that transcription factors and master growth regulators control ERES number, soma size, and dendrite size in a coordinated manner to drive neuronal cell fate determination. In this context, asymmetric cell division can be viewed as the initiating event of neuronal cell fate determination, due to uneven distribution of key resources to daughter cells.

We demonstrate in PVD that ERES number is determined by cell fate pathways rather than feedback from growing neurites. Previous work has shown that extrinsic cues such as target-derived neurotrophins regulate survival and neuronal morphogenesis demonstrating a feedback regulation mechanism (Fawcett and Keynes, 1990; Purves et al., 1988). We hypothesized that increasing secretory demand from a developing dendrite might upregulate ERES number. The PVD dendritic arbors are not only unusually large among worm neurons but also precisely target the space between skin and muscles (Zou et al., 2016). In the dendrite guidance factor and receptor mutants, the dendritic arbor is drastically smaller and fails to grow into the skin-muscle junctions (Dong et al., 2016; Liu and Shen, 2011; Zou et al., 2018, 2016). However, we found these mutants did not show abnormalities in the number of early secretory organelles in PVD. These results argue strongly that the size, shape and precise targeting of PVD dendrites are dispensable for the determination of somatic ERES number. Consistent with this notion, we demonstrate that GFP::SEC-16 puncta emerge rapidly during PVD birth, and establish high ERES numbers before neurite outgrowth. These findings suggest that secretory organelle number, and likely secretory capacity, is encoded into a neuron’s identity, acting as part of the cell-intrinsic programming that helps to drive a cell’s fate. Thus, while local neurite outgrowth is highly regulated by extrinsic growth factors and cues, overall secretory capacity, the “engine” underlying neurite growth, may be largely determined by cell fate programs driving global nutrient sensing pathways (see below).

We report that cell fate programming, governed by transcription factors UNC-86 and MEC-3, drives PVD ERES number in two phases: establishment and maintenance. In the establishment phase, our direct imaging experiments show that PVD inherits a large soma due to asymmetric cell division. We used different approaches to manipulate the size of PVD soma at birth and found that there is a strong correlation between initial cell size and ERES number. In this phase, ERESs emerge rapidly as mitosis concludes, and the establishment of ERES numbers requires both the cell fate transcription factor UNC-86 and regulators of cell division plane including LIN-5/NUMA. Asymmetric cell division has been previously shown to differentially distribute cell fate transcription factors and organelles (Li et al., 2016; Vertii et al., 2018). It is tempting to speculate that uneven distribution of ERES components, such as SEC-16 and ER membranes, could provide the necessary seeds to establish persistent differences in secretory capacities. Given that PVDsister undergoes apoptosis soon after birth, there may be a preferential sorting of pro-growth factors away from PVDsister to PVD, such that PVDsister is, in a sense, sacrificed to boost PVD’s growth potential. In the maintenance phase, MEC-3, UNC-86 and sufficient nutrients are needed to actively maintain ERES number. In *mec-3(e1338)* or *unc-86(e1416)* mutants, as well as starvation conditions, the number of ERESs decreases during this phase. The differences in ERES number phenotypes between *unc-86(e1416)* and *mec-3(e1338)* are consistent with the differential expression of these two transcription factors. Previous work shows that UNC-86 is expressed early, before PVD birth, and later activates MEC-3, which forms a heterodimer with UNC-86 to activate lineage specific transcriptional programs (Chalfie and Au, 1989; Duggan et al., 1998; Finney and Ruvkun, 1990; Xue et al., 1993, 1992). These expression patterns are consistent with our findings that UNC-86 regulates both establishment and maintenance phases of ERES number dynamics, while MEC-3 is only required for the maintenance phase. In a possible third phase, ERES numbers become stable because late starvation, in contrast to early starvation, does not lead to a reduction of ERES.

We show ceTOR/LET363 acts downstream of cell fate transcription factors during the maintenance phase, integrating nutrient availability information to maintain ERES number and promote somatodendritic growth of PVD. This work raises the possibility that cell fate transcription factors could modulate ceTOR expression levels, which, in the presence of sufficient nutrients, set biosynthetic and secretory capacities required for a particular cell fate. Such a function would indicate a new cell autonomous role for TOR as a regulator of physiological neuron cell size. TOR’s function and signaling pathways in growth and nutrient sensing has been intensively studied in various contexts, but its role in non-pathological growth of neurons *in vivo* is not well understood. Kwon *et al*. found that abnormal soma expansion in granule cells of PTEN-deficient mice was dependent on TOR (Kwon et al., 2003). Choi *et al*. showed that rapamycin-mediated inhibition of TOR in cultured embryonic rat hippocampal neurons impaired axon growth but not dendrite growth (Choi et al., 2008). We report a cell-specific requirement for *ceTOR/let-363* in the normal physiological growth of PVD. *ceTOR/let-363’s* role in PVD appears to coordinate soma size, ERES number, and dendrite size, suggesting it may be a cell-intrinsic master regulator for biosynthesis, secretion and cell size in PVD. This impact on soma size differs from the finding by Kwon *et al*. that TOR disruption did not impact WT soma size in murine granule cells (Kwon et al., 2003). Several factors may account for the apparent discrepancy in these results. Granule cells are among the smallest cells in vertebrate brains and their physiological growth may therefore be less sensitive to TOR (D’Angelo, 2016). More broadly, TOR-dependent growth may vary between neuron types and species. Further work is required to test whether differential TOR activity is a general feature of neuron size and fate regulation.

Several potential downstream mechanisms could explain the reduction of ERES number in *ceTOR/let-363* mutants and starved animals. TOR plays well-established roles in promoting protein translation and lipid biosynthesis, while inhibiting autophagy (Deleyto-Seldas and Efeyan, 2021; Keith Blackwell et al., 2019; Laplante and Sabatini, 2009; Nandagopal and Roux, 2015). It is possible that a reduction in protein translation and the resultant reduction of secretory flux subsequently reduces ERES number(Farhan et al., 2008b). In addition, a lack of TOR-mediated lipid synthesis, or a disinhibition of ER-phagy (Hamasaki et al., 2005; Zhao et al., 2015) may restrict the size of the soma and ER membrane, constraining the distance between ERES, and thus increase probability of ERES fusion to reduce ERES number(Farhan et al., 2008a; Speckner et al., 2021; Tillmann et al., 2015). Another possibility is that lack of TOR activity disinhibits autophagy of ERES directly, which may help to reduce secretion during starvation. Zacharogianni et al. show that amino acid starvation in cell lines results in a TORC1 complex-independent dispersal of ERESs (Zacharogianni et al., 2011). Together with our data suggesting a ceTOR-dependent ERES reduction, these results support the notion that secretory organelles respond to starvation via a number of different pathways depending on variables such as the particular types of nutrient perturbations or organisms under study.

The rapid emergence of GFP::SEC-16 during cell birth is reminiscent of phase transition dynamics characteristic of liquid-liquid phase separation (LLPS). Indeed, our observations of ERES establishment in PVD are consistent with several characteristics of LLPS, and suggest that regulation of biophysical properties of ERES components may play a role in determining ERES number. For instance, the dependence of initial ERES number on PVD cell size is consistent with the inherent size scaling behavior of LLPS condensates (Brangwynne, 2013). The tools developed here enable additional studies to further assess the emerging role of LLPS in the establishment and maintenance of ERESs in multicellular organisms (Alberti et al., 2019; Bevis et al., 2002; Farhan et al., 2008a; Gallo et al., 2020; Heinzer et al., 2008; Peotter et al., 2019; Speckner et al., 2021; Tillmann et al., 2015).

Taken together our data suggest a model for neuronal cell size control. Cell fate transcription factors in PVD drive asymmetric cell division to establish a large initial cell size, which, in turn, determines high ERES number, perhaps through LLPS size scaling. ERES number is subsequently maintained via a transcription factor-mediated modification of TOR levels, which regulates the nutrient-dependent biosynthetic and secretory capacity required for PVD’s neuronal fate. Our data also provide an additional perspective for future investigation into why neurons are particularly sensitive to mutations in early secretory pathway proteins (Tang, 2021). The upregulation of early secretory pathway structures shown here suggests that larger neurons may be particularly reliant on high, efficient secretory output. Large post-mitotic neurons require a high minimum secretory output to establish and maintain the expansive morphologies critical to their function. Thus, relatively small defects in secretory capacity may disproportionately impact form and function of large neurons. Thus, investigation into how large neurons regulate secretory structure and function will offer valuable insights into the heightened vulnerability of nervous systems to early secretory pathway defects.

## Methods

### *C. elegans* Strains

All strains were grown on nematode growth medium (NGM) plates seeded with OP50 *E. coli* at 20°C unless otherwise stated for experimental conditions. N2 Bristol was used as wild-type strain. Mutant alleles and transgenes used in this study are listed in Table S1. *lin-5* mutants were kindly provided by Dr. Xing Liang.

### Plasmids

Plasmid constructs for *C. elegans* expression were generated in pSM delta vector as a backbone, which was kindly provided by Dr. Andrew Fire. KE51(*Pmec-17*::mScarlet) & *Plin-32::FLP*, and pCER218 (*ser2prom3:: sar-1* (T35N)) constructs were kindly provided by Dr. Kelsie Eichel and Dr. Claire Richardson. DNA constructs were assembled into plasmid constructs using Gibson cloning methods. pCER218 (Pser2prom3:: *sar-1* (T35N), called *sar-1DN* in text) was made by amplifying the worm genomic *sar-1*, cloning it into the pSM delta vector and introducing a T35N mutation in a highly conserved residue in the GTPase domain (Kuge et al., 1994) to make *sar-1* (T35N) via site-directed mutagenesis.

### *C. elegans* transformation

Transgenic extrachromosomal arrays were generated using a standard microinjection protocol (Mello and Fire, 1995). In short, plasmid constructs and co-injection marker were mixed and injected to distal arm of the gonad of young adult animals. F1 animals were then screened for transgenic lines. The details of transgenic arrays can be found on Table S1.

### CRISPR Genome editing

CRISPR knock-ins were generated by direct gonadal injection of Cas9 protein (IDT) duplexed with tracrRNA and targeting crRNA (IDT) along with respective repair templates (Paix et al., 2017). For long insertions (> 150bp), knock-in templates (GFP-FLPon-SEC-16A.2) were PCR amplified from pSK–FLPon– GFP vectors (McDonald et al., 2020) with ultramer oligonucleotides (Integrated DNA Technologies) containing 100 bp of flanking genome homology. PCR products were purified by ethanol precipitation or PCR cleanup and used for injection at 300ng/μl concentrations. For short insertions (loxp sites), synthetic single-stranded DNA oligos (IDT Ultramers) were used as templates and injected at 600nM concentrations. F1 animals were screened by PCR genotyping. Heterozygous F1 animals were homozygoused and the insertions were verified by Sanger sequencing. The following guide RNA sequences were used for genome edits: GFP-FLPon-SEC-16A.2: TGTTGCCAATAGAAGCTCAT; *let-363*-loxp: tttttctatattcgaaccaa & tgatattttgaaatatgcaa; *rab-1*-loxp: gaggaatcgatgcgagaatg & gtgaaagagtgccgaagagg.

### *C. elegans* synchronization, staging and nutrient deprivation

Gravid adults were washed off from eight to ten 6cm-plates with M9 buffer. Animals were then bleached in hypochlorite solution to obtain embryos. Embryos were washed with M9 buffer two to three times and then resuspended in M9 and allowed to hatch overnight into L1-arrested animals. L1-arrest animals were kept for maximum of 24hr before plating onto OP50-seeded NGM plates.

For the nutrient deprivation protocol, arrested L1s were plated onto freshly seeded OP50 NGM plates, with varying amounts of OP50 in order to vary developmental stage of starvation. Animals were then monitored over the next 20-30 hr, and at the appropriate stage (either several hours before PVD-birth or immediately following PVD-birth) worms were either allowed to continue growing on starved plates or removed from OP50 onto either nutrient-free NGM plates (for starvation), or OP50-seeded NGM plates (fed controls). Data from fed and starved worms was collected the following day.

For time course experiments, animals were synchronized as described above and examined under a Zeiss Axioplan fluorescence scope with 63X Plan-Apochromat 63 × /1.4 NA objective for counting of ERES at indicated timepoints.

To image early PVD morphogenesis events (L2 animals), L1-arrest animals were grown on OP50-seeded NGM plates for 25-28 hr depending on the strain at ~20°C to reach L2 stage. After L2 stage has been reached, animals were monitored every 30 minutes under Zeiss Axioplan fluorescene scope with 63X Plan-Apochromat 63 × /1.4 NA objective. Animals were mounted on slides for imaging when majority of animals reached desired developmental stage (PVD precursor stage). To image mature PVDs and PDEs and TRNs, L4 animals, identified based on vulval development, were mounted on slides.

### Microscopy

For still confocal images, hermaphrodite worms at L2 or L4 stage worms were immobilized on 2-3% agarose pads using 10mM levamisole in M9 buffer. Images were obtained on a spinning disk system (3i) with a CSU-W1 spinning disk (Yokogawa), 405-nm, 488-nm and 561-nm solid-state lasers, a C-Apochromat 63×/1.2 NA water-immersion objective, and a Prime95B camera (Photometrics). Z-stacks with 0.25-0.5 μm sections spanning the imaged neurons were taken. For live-imaging of developing worms, L2 stage hermaphrodite worms were immobilized on 7.5% agarose pads using 2.5mM levamisole in M9 buffer. Slides were sealed with VALAP (Vaseline, lanolin, paraffin, 1:1:1) to prevent agarose pad from desiccation. FRAP of ERES was performed on the same system with a Vector diffraction-limited xy scanner (3i). Live-imaging was performed with the same system. Z-stacks with 0.4 μm sections were obtained every 3 minutes for maximum duration of 6 hours.

### Image Analysis

ERES were counted manually from unmodified z-stacked images, using the neuron plasma membrane marker mCherry::PH(PLCγ), or cytoplasmic mScarlet to identify cells of interest (COI), and counting the number of endogenous GFP::SEC-16A.2 puncta in COI somas. Two puncta that appeared to be in contact but had clearly separate centers of highest intensity were defined as doublets and counted as two ERESs. Counting criteria were the same for counts made in z-stack images taken on the 3i spinning disk confocal and counts made on animals directly under the Zeiss Axioplan microscope under the same magnification. For fluorescence intensity quantifications of nuclear-localized Plet-363::GFP, total fluorescence intensity values were obtained at ROIs of SUM-projected Z-stacks of PVD and PDE nuclei using ImageJ.

### Electron microscopy

Wild-type N2 worms were prepared for conventional EM by high pressure freezing/freeze-substitution. Worms in M9 containing 20% BSA and *E. coli* were frozen in 100 μm well specimen carriers (Type A) opposite a hexadecane coated flat carrier (Type B) using a BalTec HPM 01 high-pressure freezer (BalTec, Lichtenstein). Freeze-substitution in 1% OsO_4_, 0.1% uranyl acetate, 1% methanol in acetone, containing 3% water (Buser and Walther, 2008; Walther and Ziegler, 2002) was carried out with a Leica AFS2 unit. Following substitution, samples were rinsed in acetone, infiltrated and then polymerized in Eponate 12 resin (Ted Pella, Inc, Redding, CA). Serial 50 nm sections were cut with a Leica UCT ultramicrotome using a Diatome diamond knife, picked up on Pioloform coated slot grids and stained with uranyl acetate and Sato’s lead (Sato, 1968). Sections were imaged with an FEI Tecnai T12 TEM at 120 kV using a Gatan 4k x 4 k camera. TrakEM2 in Fiji was used to align serial sections (Cardona et al., 2012; Saalfeld et al., 2012; Schindelin et al., 2012). Modeling of serial sections was performed with IMOD (Kremer et al., 1996).

### Neuron Soma Measurements

Z-stack still images taken on the 3i spinning disc microscope, as described above, were projected into a single plane using ImageJ software. ROI’s were drawn around the projected soma, and the area of this ROI was measured in μm^2^ as an estimation of the maximal cross-section of the cell in the xy plane.

### Quantification of dendritic arbor

For Figure 1c, all terminal branches in AVM, PVM, PDE, and FLP were counted in images of L4 animals. For all PVD dendrite quantifications, images were straightened with the ImageJ software (US National Institutes of Health) using PVD primary dendrite as a reference. For WT PVD in Figure 1c, quaternary branches were counted within a rectangular region of interest (ROI) extending 150μm anterior of PVD soma and 30μm posterior of PVD soma. For SAR-1-DN and control L4 animals (Fig. 1i), a 100μm long rectangular ROI was placed starting at the center of PVD soma extending 100 μm anteriorly, containing the entire width of the animal, while for *rab-1(wy1375)* (Fig. 1h), SAR-1DN, and control young adult animals, a 100μm long rectangular ROI was placed with PVD soma at the center, and the number of secondary, tertiary and quaternary branches within ROIs were counted. For *let-363*-floxed and control L4 animals, all branches anterior of PVD soma were counted. Any protrusion from the primary dendrite was classified as a secondary branch, and any correctly-oriented protrusion from the tertiary dendrite was classified as a quaternary branch.

### Quantification and statistical analysis

Quantifications were blinded in any cases where mutant phenotypes were not easily observable. All statistical analysis was done using GraphPad Prism 9 software. All statistical parameters can be found in the corresponding figure legends. No statistical methods were used to predetermine the sample size.

### Materials and data availability

All reagents and data are available upon request.

## Acknowledgements

We thank C. Wicky and F. Muller for strain FR297 (Long et al., 2002); X. Liang for *lin-5* reagents; C. Tzeng for expression constructs; J. Lee for assistance with long-term time-lapse imaging; L. Tao for guidance receptor mutant morphology images; C. Richardson and members of the Shen Lab for conversation and reagents.

---

**Fig. S1.**
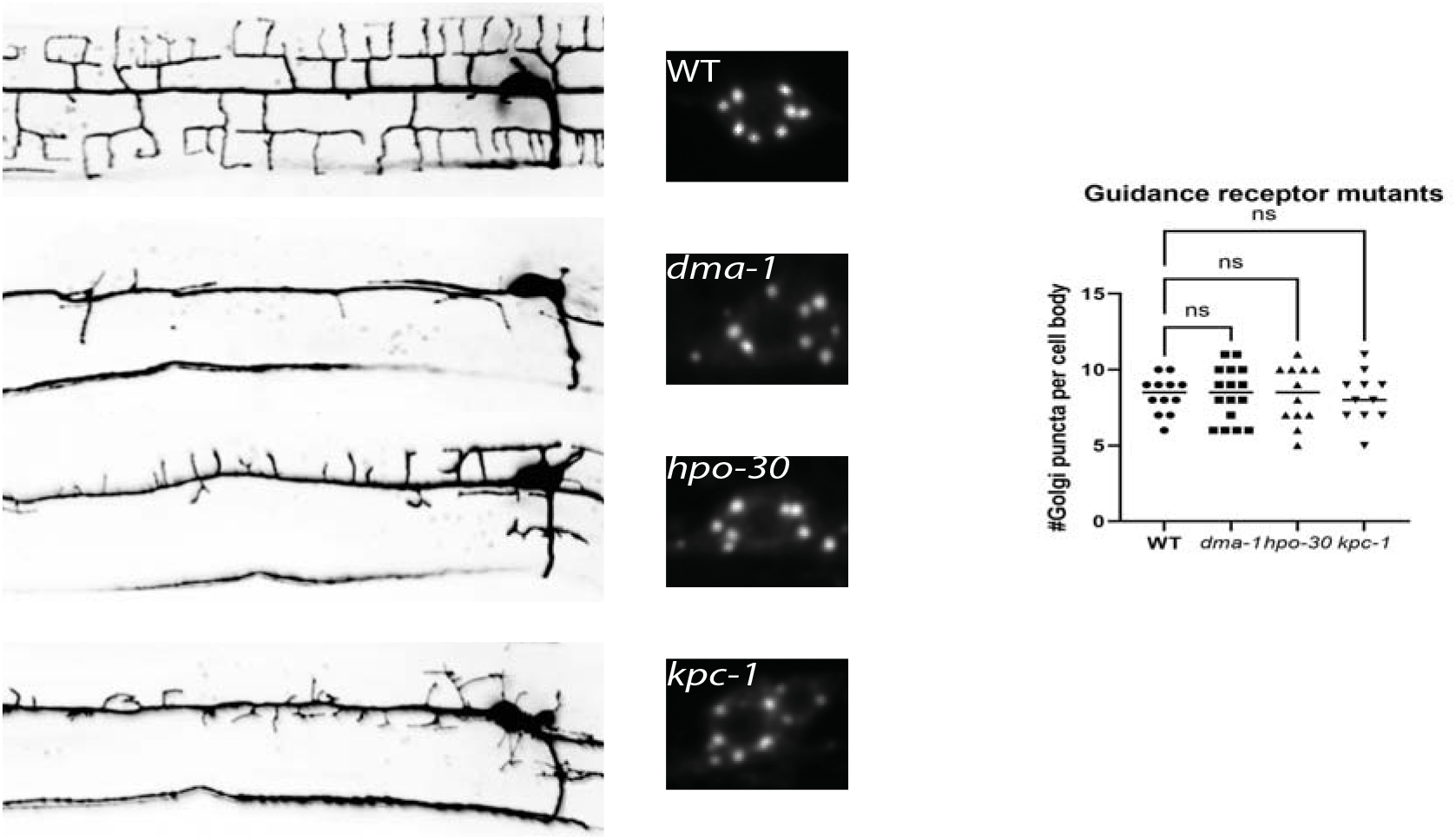
(related to Fig. 2): Guidance receptor mutants do not impact the number of early secretory structures. Left: PVD dendrite morphology of wild type and indicated mutants. Middle: AMAN-2::G-FP overexpression shown in PVD soma. Right: quantification of number of AMAN-2::GFP puncta in somas of wild type and indicated mutants. See methods Table S1 for allele names.

**Table S1:**
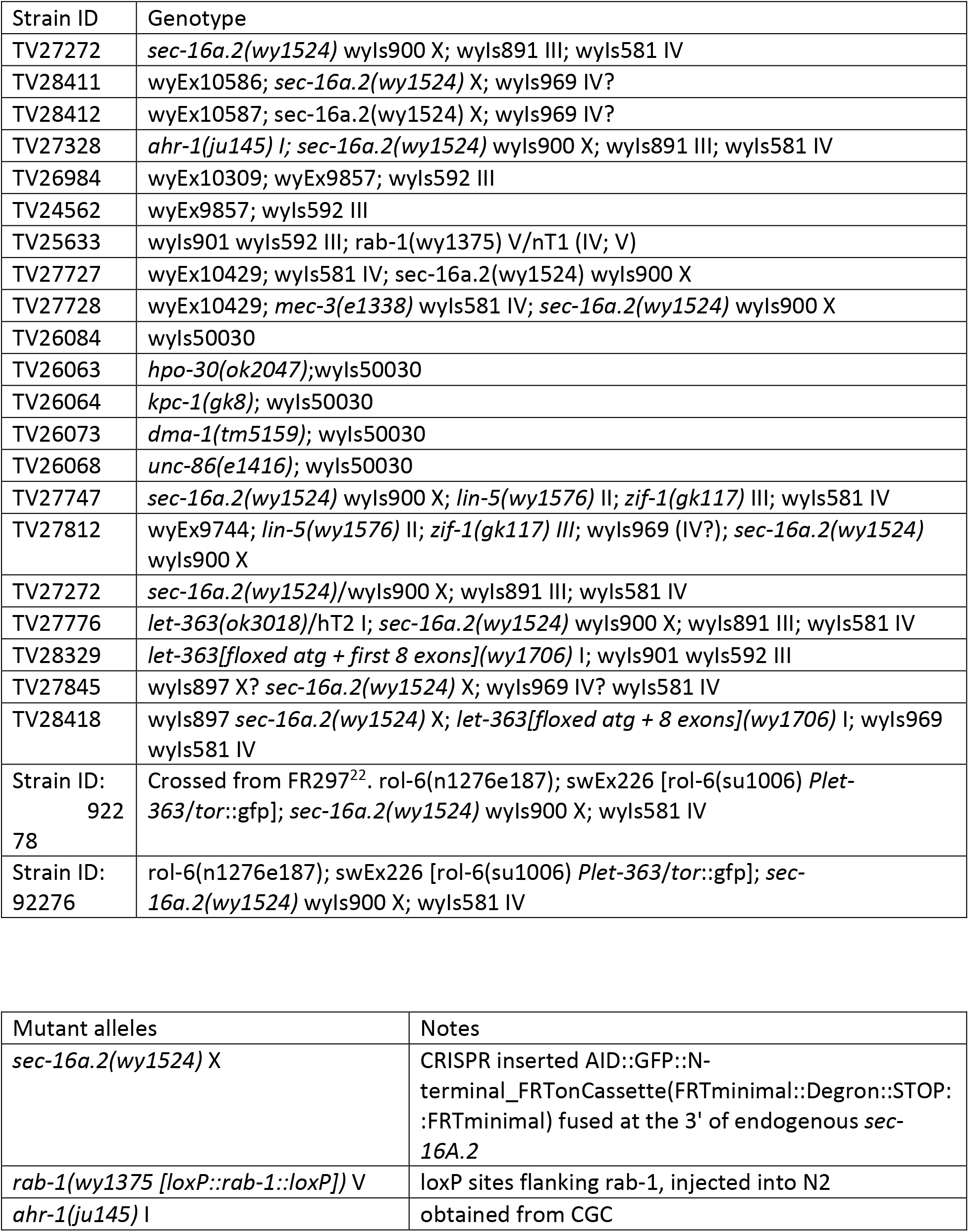
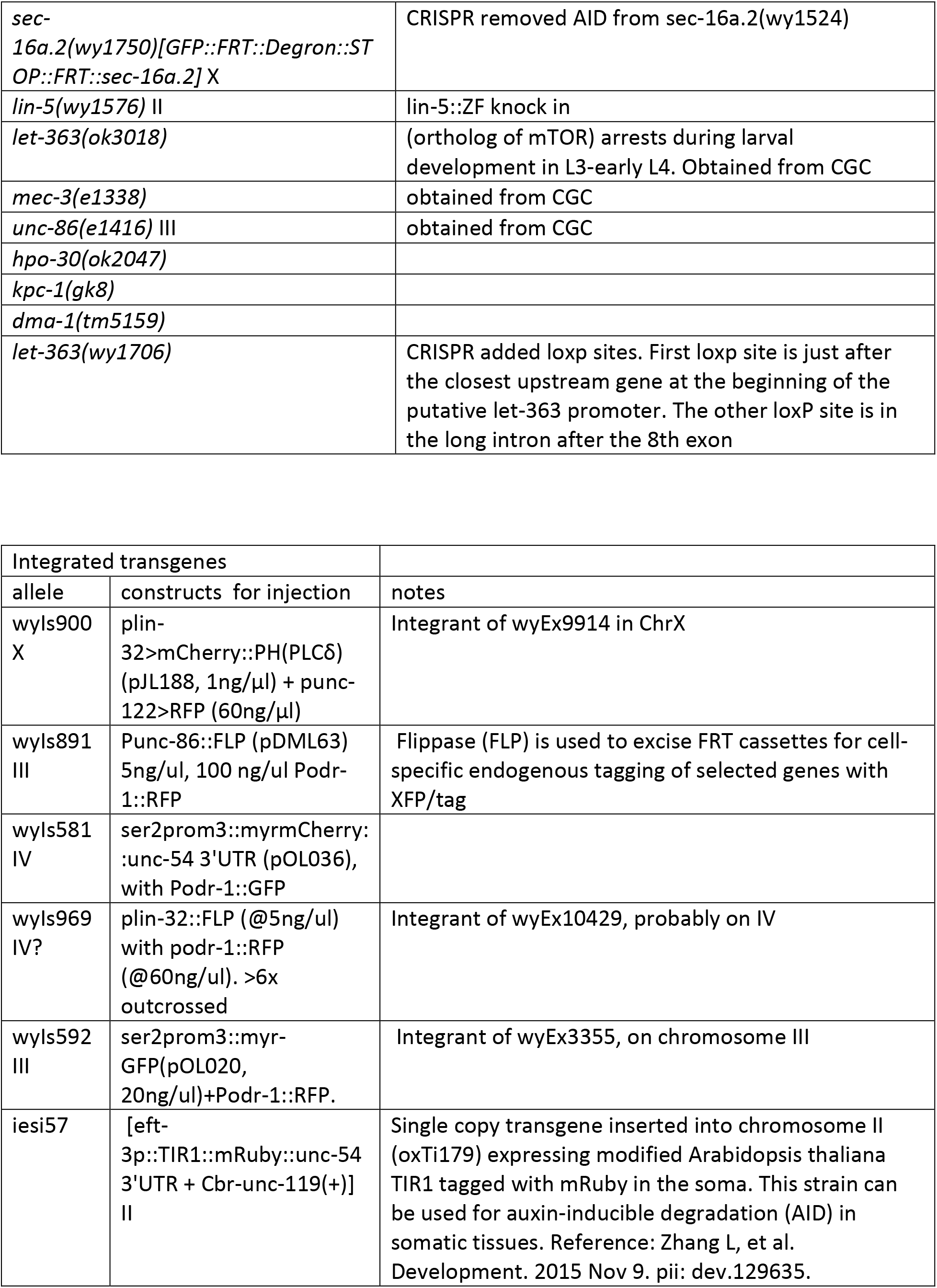
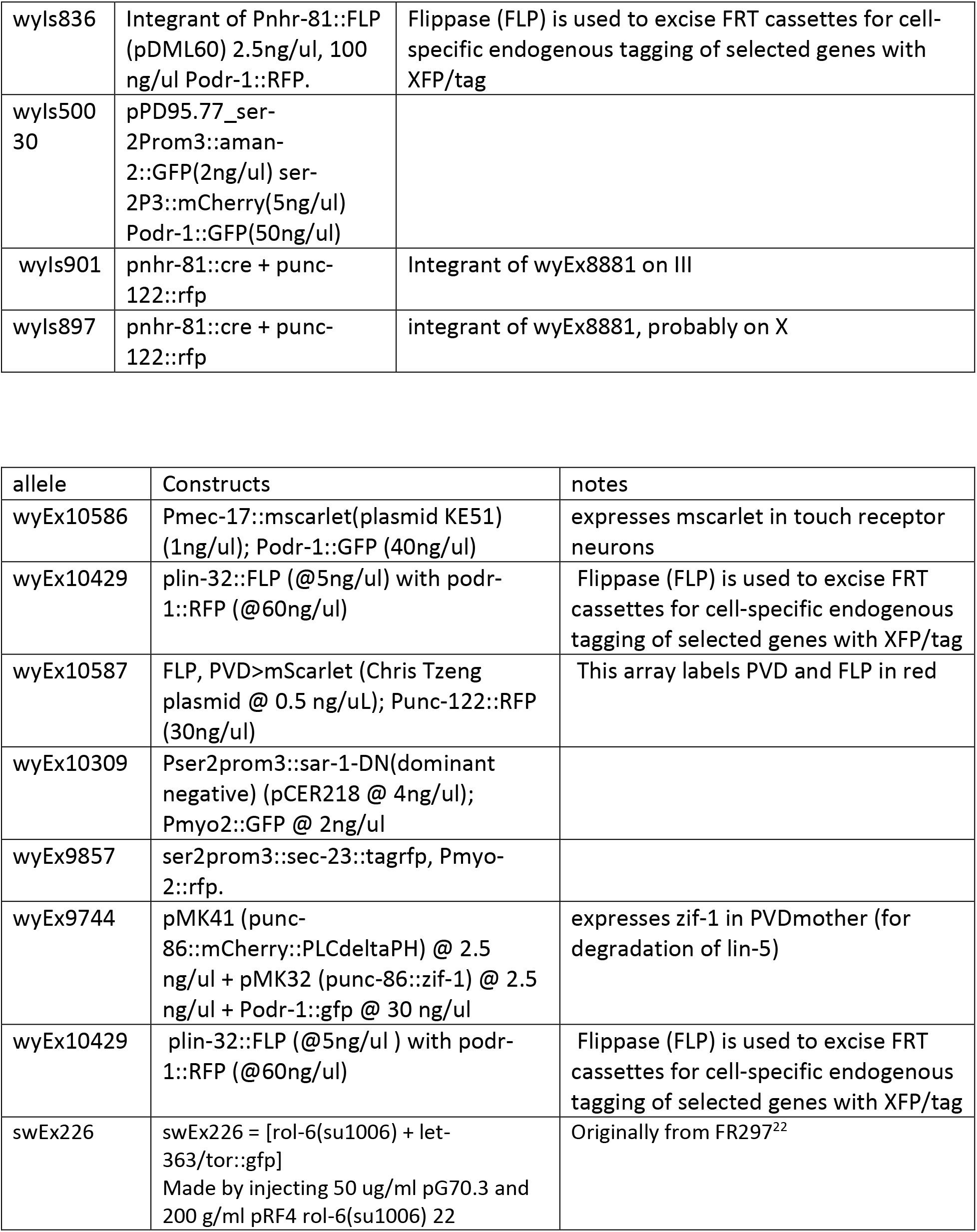

