## Supplemental Figure S1 for "Neurons alter endoplasmic reticulum exit sites to accommodate dendritic arbor size"

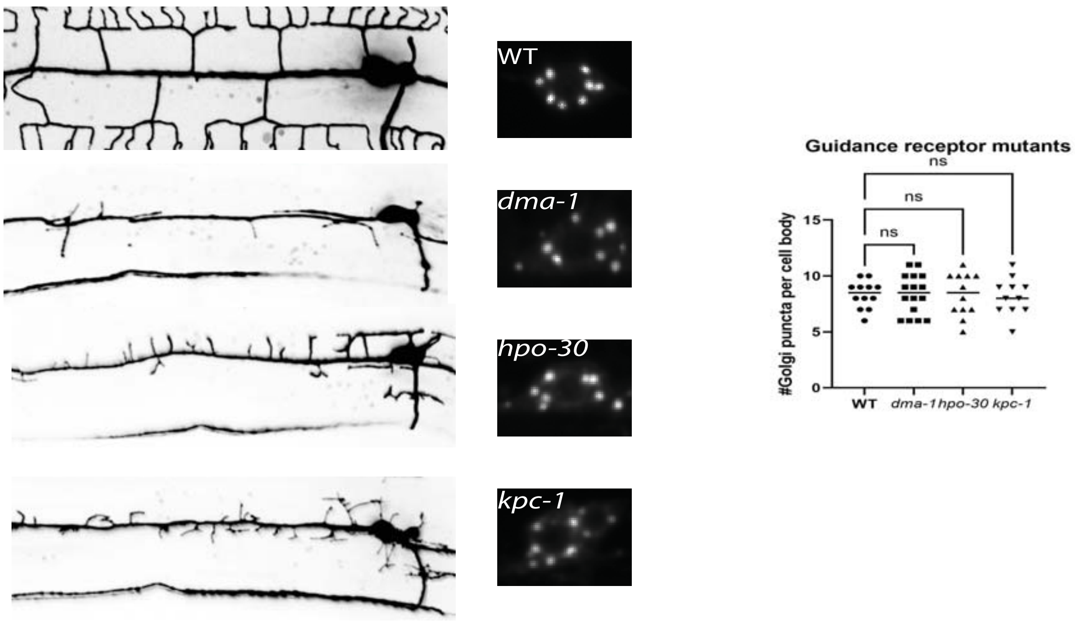


Fig. S1 (related to Fig. 3): Guidance receptor mutants do not impact the number of early secretory structures. Left: PVD dendrite morphology of wildtype and indicated mutants. Middle: AMAN-2::GFP overexpression shown in PVD soma. Right: quantification of number of AMAN-2::GFP puncta in somas of wildtype and indicated mutants. See methods Table S1 for allele names.
